# Synergistic antifungal effect of amphotericin B-loaded PLGA nanoparticles based ultrasound against *C. albicans* biofilms

**DOI:** 10.1101/423723

**Authors:** Min Yang, Kaiyue Du, Yuru Hou, Shuang Xie, Yu Dong, Dairong Li, Yonghong Du

## Abstract

*C. albicans* is human opportunistic pathogens that cause superficial and life-threatening infections. An important reason for the failure of current antifungal drugs is related to biofilm formation mostly associated with implanted medical device. The present study aims to investigate the synergistic antifungal efficacy of low-frequency and low-intensity ultrasound combined with amphotericin B-loaded PLGA nanoparticles (AmB-NPs) on C.albicans biofilms. AmB-NPs were prepared by a double emulsion method and demonstrated the lower toxicity than free AmB, after which biofilms were established and treated with ultrasound and AmB-NPs separately or jointly in vitro and in vivo. The results demonstrated the activity, biomass, and proteinase and phospholipase activities of biofilms were decreased significantly after the combination treatment of AmB-NPs with 42 KHz ultrasound irradiation at an intensity of 0.30 W/cm^2^ for 15 min compared to the control, the AmB alone or the ultrasound alone treatment (P < 0.01), and the morphology of biofilms was altered remarkably after jointly treatment under CLSM observation and detection, especially thickness thinning and structure loosing. Furthermore, the same synergistic effects were proved in a subcutaneous catheter biofilm rat model. The result of colony forming units of catheter fungus loading exhibited a significant reduction after AmB-NPs and ultrasound jointly treatment for 7 days continuous therapy, and the CLSM images revealed that the biofilm on the catheter surface was substantially eliminated. Our study may provide a new noninvasive, safe and effective application to C.albicans biofilm infection therapy.

## Introduction

*C. albicans* is a normal commensal organism found on human mucosal surfaces that acts as an opportunistic pathogen in immunocompromised patients. In past decades, the infection rate and mortality rate of *C. albicans* infections has been increasing annually in response to the widespread use of antibiotics and hormone treatments. Approximately 90% of *C. albicans* infections are related to biofilm formation associated with medical devices, such as indwelling catheters, venous catheters for parenteral nutrition, prosthetic valves, and joint implants^[1-2]^. Biofilms are communities of microorganisms that attach to the surface of tissues or biological materials and encase themselves in self-secreted extracellular matrix (EMC, major sugars and proteins), which form a dense natural barrier to evade the immune function of the host and prevent permeation of antibiotics^[3]^. Biofilm formation is an important reason for the failure of current antifungal drugs because it makes microbes less susceptible to antifungal agents, imparting 10–2000 times higher resistance to the effects of antimicrobial agents^[4-5]^. Currently, there are only few antimycotics, including azoles (miconazole), polyenes (amphotericin B, AmB) and echinocandins, which are partially effective against biofilm-associated infections. However, these sometimes cause significant adverse effects when applied via parenteral administration, such as nephrotoxicity hepatotoxicity, hematotoxicity, and hypersensitivity reactions, which has limited its widespread use^[6-7]^. Therefore, the newer antifungal drugs with greater hypotoxicity and greater efficacy, as well as newer treatment methods that can reduce the amount of drugs and achieve effective antifungal effects, have become a focus of studies investigating anti-biofilm infections.

Naturally degradable and synthetic biodegradable polymer nanomaterials for alternative antibacterial therapy have been extensively reported as innovative tools for combating the high rates of antimicrobial resistance, including multidrug-resistant (MDR) bacterial and biofilm-associated infections^®^. Among them, poly(lactic-co-glycolic acid) (PLGA) biodegradable polymeric nanoparticles-based drug delivery systems have been proposed as a potential alternative to various biomedical applications for vaccination, cancer, inflammation and other diseases^[9-10]^. Moreover, oral or parenteral administration of AmB polymeric nanoparticle formulations have been reported to have antifungal efficacy and lower toxicity with increased bioavailability compared to intravenous Ambisome^®^ or Fungizone^™^ ^[11-12]^. Thus, we investigated the ability of synthesized AmB-loaded PLGA nanoparticles (AmB-NPs) to reduce the adverse effects of free AmB and explored the antifungal efficacy of AmB-NPs toward *C. albicans* biofilms in this study. During biofilm formation, surface-adherent extracellular matrices of the biofilm act as a barrier to antibiotic diffusion, which limits the exposure of intracellular bacteria in the biofilms to superficial antibiotics. Ultrasound has been widely acknowledged as a promising approach to overcome these obstacles by enabling increased permeability for gene/drug delivery^[13-14]^. Ultrasound exposure induces sonoporation, which may transiently disrupt the cell membrane and increase the permeability of the membrane to promote transportation of membrane-impermeable agents, such as nanoparticles or antibodies, into cells or tissues in preclinical studies. Ultrasound radiation also creates many holes in the extracellular matrix of bacterial biofilms, which can benefit the rapid uptake of bulk antibiotics, oxygen and nutrition by cells ^[15-16]^. More importantly, the presence of nanoparticles can enhance acoustic cavitation by increasing cavitation nuclei to reduce the initial energy of the cavitation threshold^[17-18^]. Our previous study also revealed that low-frequency and low-intensity ultrasound combined with AmB-loaded PLGA nanoparticles (AmB-NPs) has higher antifungal efficacy than free AmB alone against planktonic *C. albicans* in vitro^[19]^.

Therefore, we investigated the effects of low-frequency and low-intensity ultrasound on biofilm and antifungal efficiency of ultrasound combined with AmB-NPs against *C. albicans* biofilms in vitro. We also investigated the synergistic antifungal efficacy in vivo using a rat model of *C. albicans* biofilms-associated catheter infection. The results presented herein provide a new noninvasive, safe and effective treatment for *C. albicans* biofilm infection therapy.

## Results

### NP characteristics and drug release

The TEM photomicrographs demonstrated that a majority of nanoparticles were spherical, in the nanometric range, with smooth surfaces and favorable dispersibility. When compared with blank NPs, AmB-NPs clearly revealed that the drug was loaded in nanoparticles (Fig. 2A). The characteristics of D, ZP, PDI, LC% and EE% of NPs and AmB-NPs are summarized in Table 1, which showed that the drug loading content reached 5.7% and the encapsulation efficiency reached 85%.

**Fig.1.**
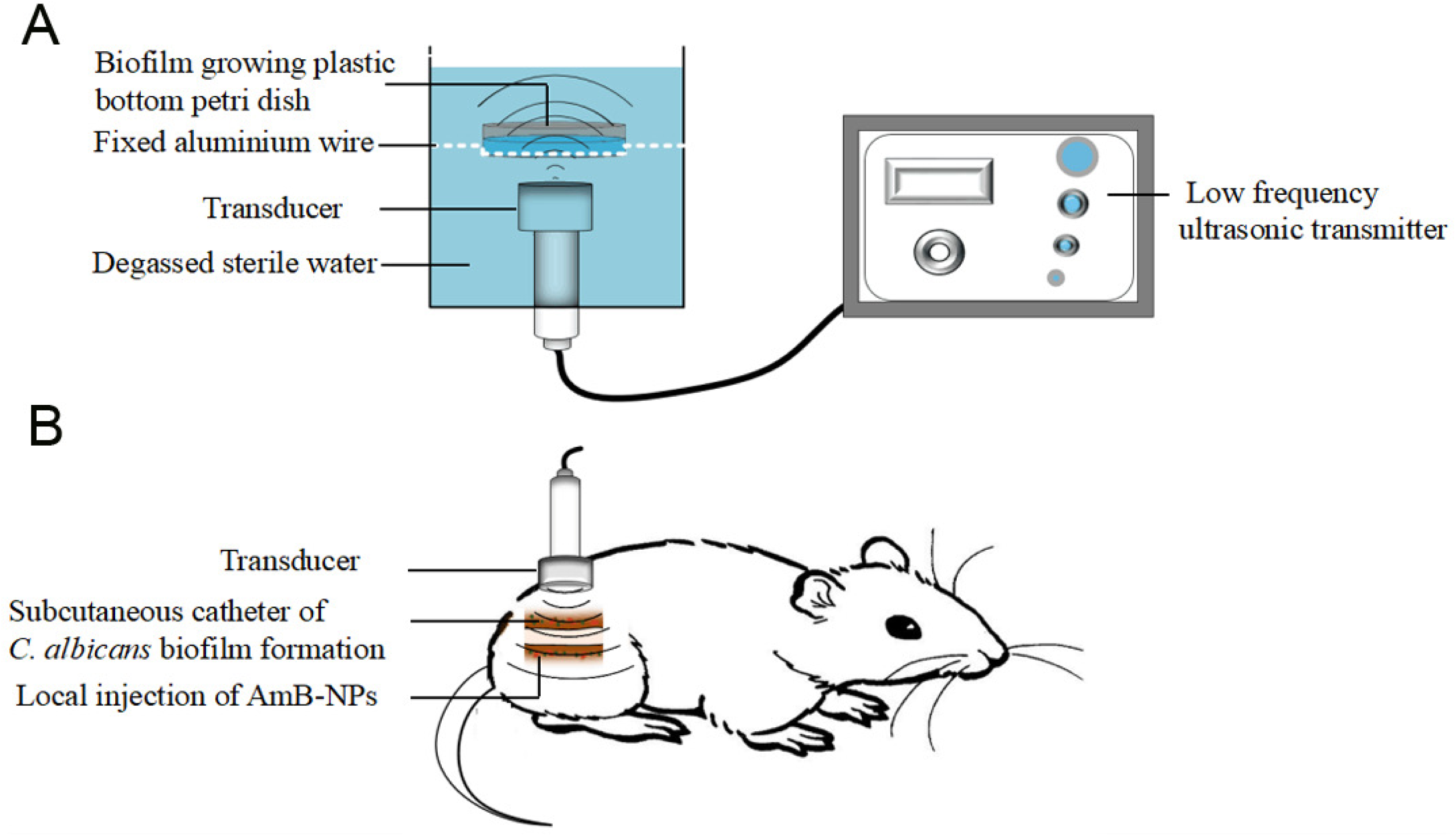
Schematic illustration of low-frequency ultrasound radiation method in vitro (A) and in vivo (B).

**Fig. 2.**
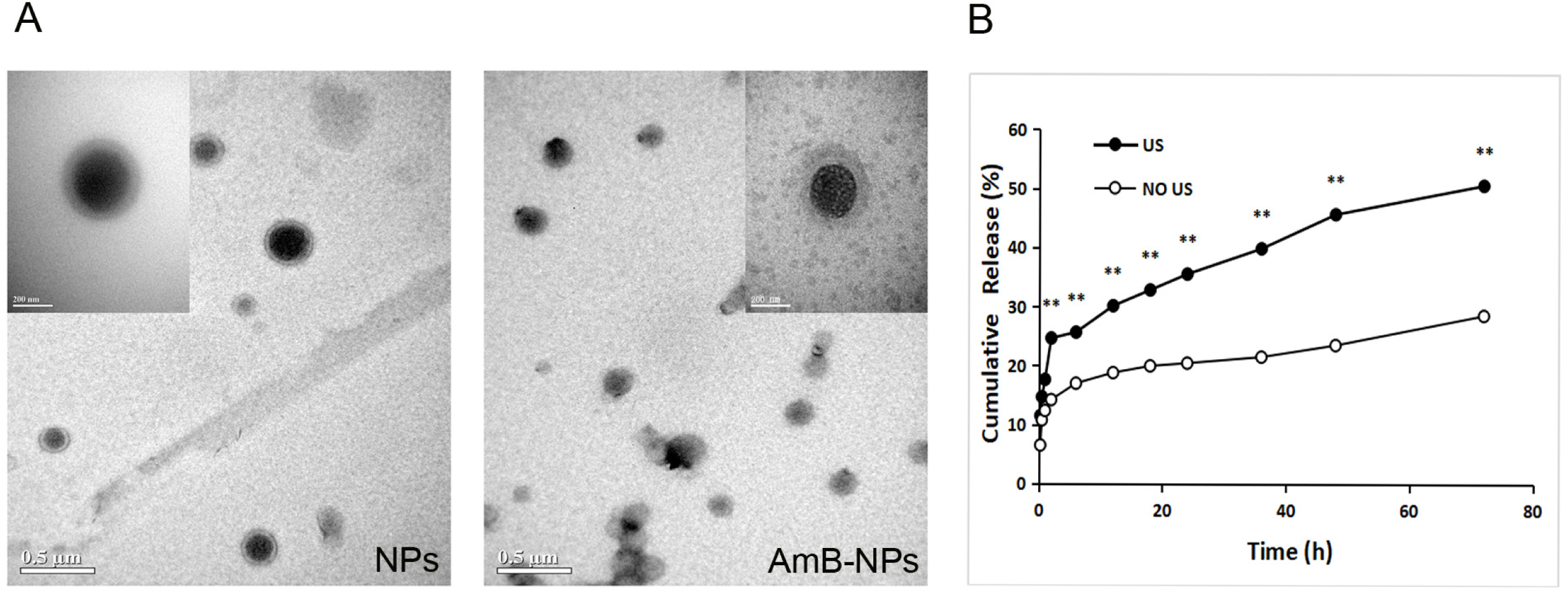
TEM photomicrographs of NPs and AmB-NPs at 20,000× or 50,000× magnification (A). The percentage of AmB cumulative release from nanoparticles with and without sonication during 0-72 h incubation (B). The double asterisk (^∗∗^) denotes a significant difference (P <0.01) between with and without sonication. NPs: plain PLGA nanoparticles; AmB-NPs: amphotericin B-loaded nanoparticles; US: ultrasound

**Table 1.**
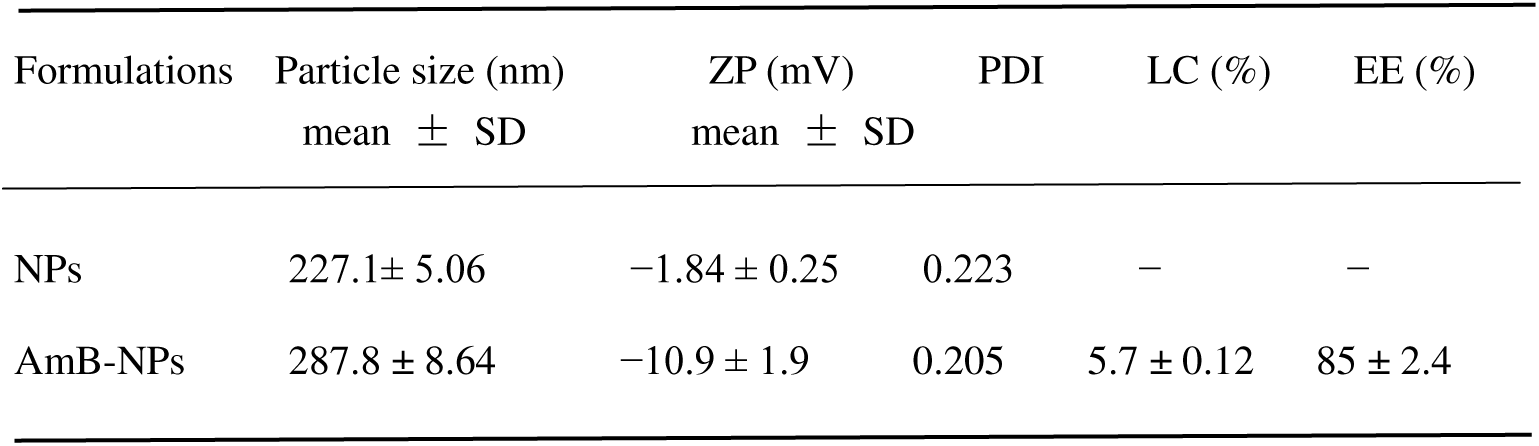
The physical characteristics of nanoparticle formulations.

The kinetic release of AmB from nanoparticles in vitro with ultrasonication or natural release is shown in Fig. 2B. The results revealed an initial burst release of about 15% of AmB in the first 2 h, followed by a continuous slow release of the drug over 72 h. Moreover, a significant higher accumulative concentration of AmB was observed after ultrasonic irradiation compared to the natural release over the same time frame. After 72 h, drug release following ultrasonic irradiation was twice as likely as natural release.

### In vitro hemolytic activity and macrophage viability of AmB-NPs

The hematological toxic reactions on red blood cells, hemolytic activity and macrophage viability of AmB-NPs relative to free AmB are depicted in Fig. 3. In the hemolytic assay with red blood cells, free AmB induced higher hemolytic activity (65.81%) than AmB-NPs (30.24%) at the same concentration of 32 μg/mL of AmB, and AmB was more likely to be hemolytic, even at very low concentrations. Moreover, NPs showed negligible hemolysis, even with increased concentrations of NPs (Fig.3A). The viability of macrophages was also significantly decreased after 24 h of co-incubation with free AmB compared to AmB-NPs at the same concentration of 4.0 μg/mL of AmB (P <0.05) (Fig.3B). These results support that AmB-NPs has better bio-safety than free AmB.

**Fig. 3.**
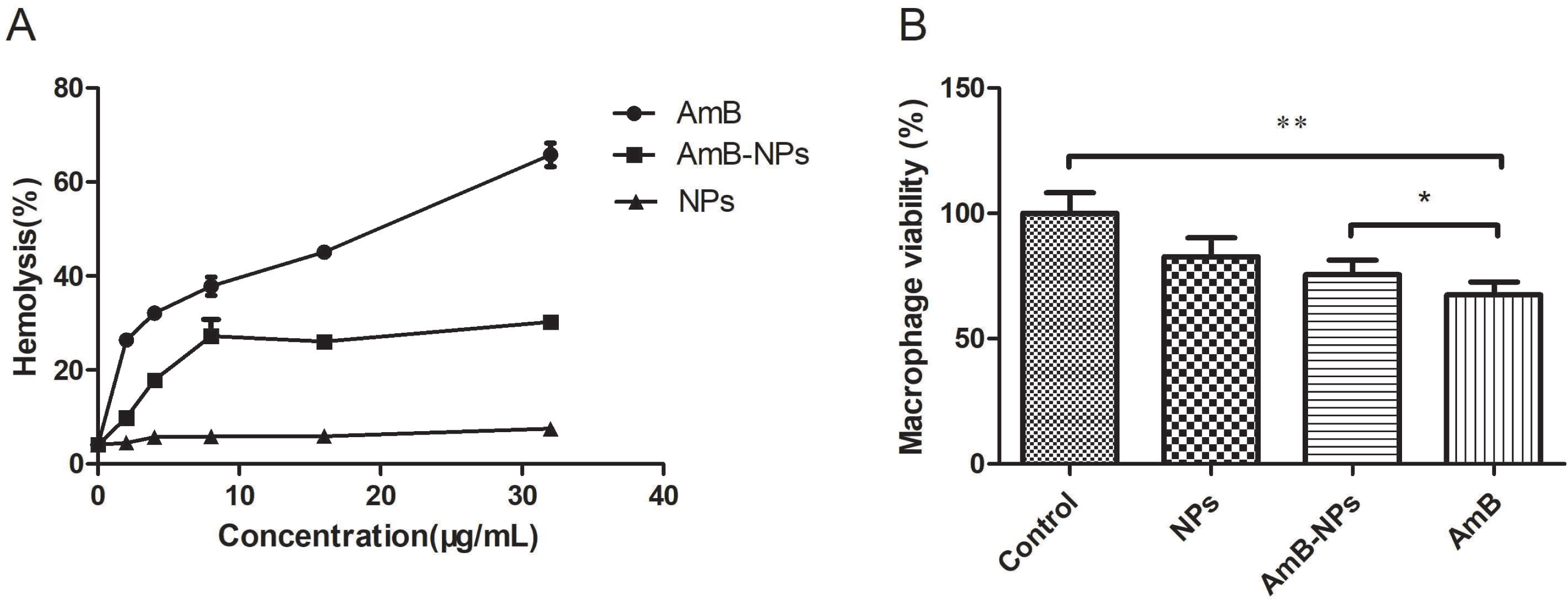
Hemolysis of red blood cells following incubation with free AmB and AmB-NPs at final AmB concentrations of 0–32 μg/mL(A). Toxicity analysis of macrophages viability following incubation with free AmB and AmB-NPs at the same AmB concentration of 4.0 μg/mL (B). ^∗∗^p<0.01 when compared with the controls, ^∗^p<0.05 when compared with the free AmB treatment. AmB: amphotericin B; NPs: plain PLGA nanoparticles; AmB-NPs: amphotericin B-loaded PLGA nanoparticles.

### Biofilm activity following a combination of ultrasound and AmB-NP treatment

To investigate the synergistically antimicrobial effect of ultrasound combined with AmB-NPs on *C. albicans* biofilm, the activity of biofilm and biofilm biomass was quantified after different treatments and the results were described in Fig. 4. The activity of biofilm significantly declined in the group jointly treated with ultrasound irradiation and 4.0 μg/mL of AmB-NPs compared to the control group, the AmB alone group or the ultrasound combined with free AmB group (P < 0.01) (Fig.4A). Additionally, the total biofilm biomass of the group jointly treated with ultrasound and AmB-NPs was also significantly lower than that of the control group (P < 0.01) (Fig.4B). However, the reduction in biofilm biomass was less than the decrease in biofilm activity, which suggested that the joint treatment with ultrasound and AmB-NPs primarily inhibited the activity of fungal cells within the biofilm.

**Fig. 4.**
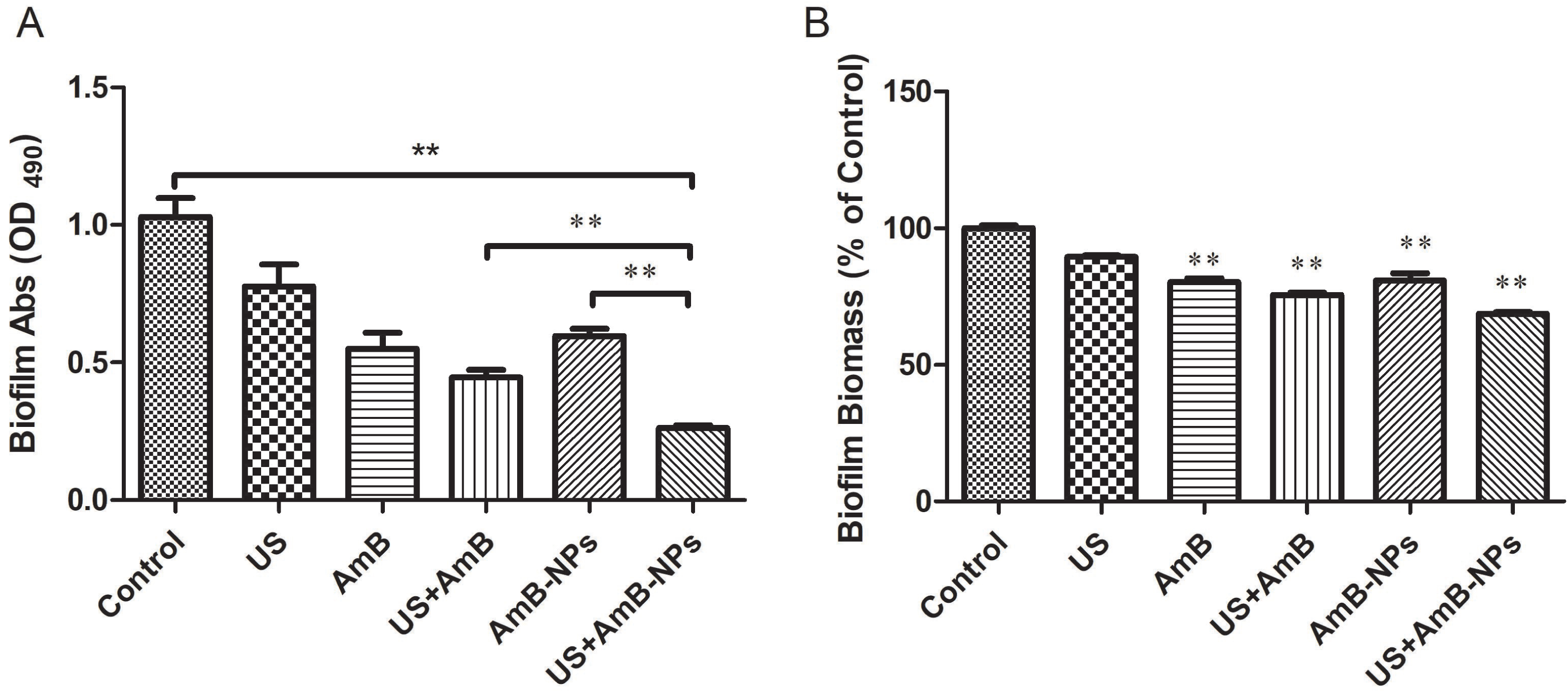
The activity of biofilm (A) and the biofilm biomass (B) all decreased significantly following treatment with a combination of ultrasound and AmB-NP compared with the control group. Ultrasonic irradiation parameters with a sound intensity of 0.30 W/cm^2^ for 15 min were chosen, and the final concentration of AmB in group AmB and AmB-NPs was 4.0 μg/mL. The double asterisk (^∗∗^) denotes a significant difference (P < 0.01) compared with the control group. US: ultrasound; AmB: amphotericin B; AmB-NPs: amphotericin B-loaded PLGA nanoparticles.

### CLSM image and COMSTAT analysis of biofilms

Biofilm CLSM images were used to observe changes in the architecture of biofilms and the living and dead fungi in different layers of the biofilm in different treatments. Figure 5 shows the viable cells (green-fluorescent) and dead cells (red-fluorescent) in the biofilm middle layer of 3-D reconstructed images. The control group was dominated by green fluorescence, which showed dense growth of live fungal cells. However, after ultrasound irradiation with AmB-NPs treatment, biofilm mainly consisted of red fluorescence (dead cells), and was characterized by sparse structures of voids, channels, pores, reduced thickness, and basically no remaining active biofilm. To further quantify the different parameters of biofilm architecture changes, the COMSTAT software was applied to analyze the mean thickness (μg), textural entropy, areal porosity and average diffusion distance (Table 2). Following ultrasound irradiation with AmB-NPs treatment, the average thickness of the biofilm was reduced by more than half, TE and ADD significantly decreased and AP markedly increased compared with the control group. Taken together, these findings confirmed that ultrasonic treatment combined with AmB-NPs has caused obvious damage to the biofilm structure of *C. albicans* in vitro.

**Fig. 5.**
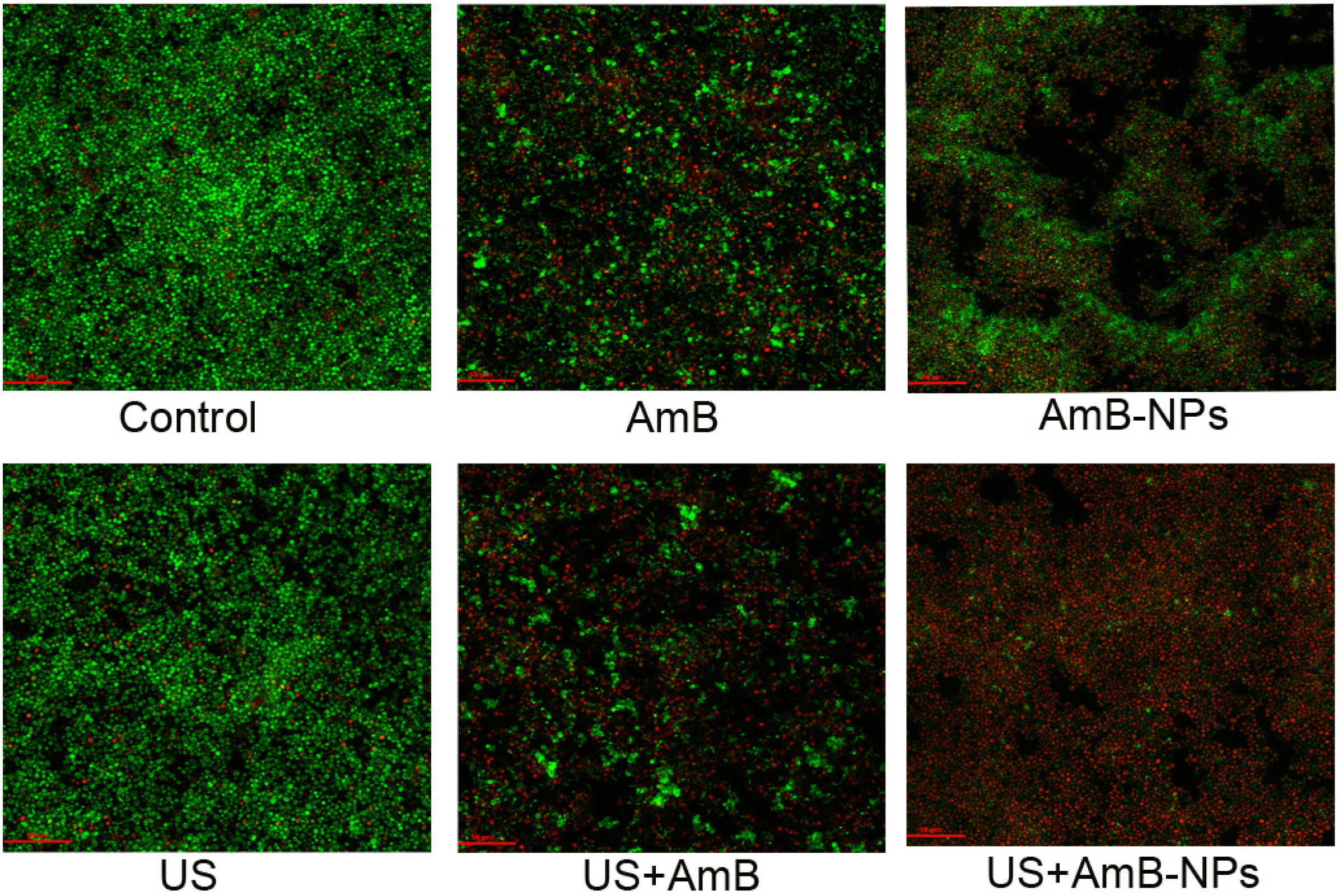
Living and dead fungal cells of the biofilm middle layer of 3-D reconstructed CLSM images in different treatment groups at 400× magnification (green, live cells)/PI (red, dead cells).

**Table 2.**
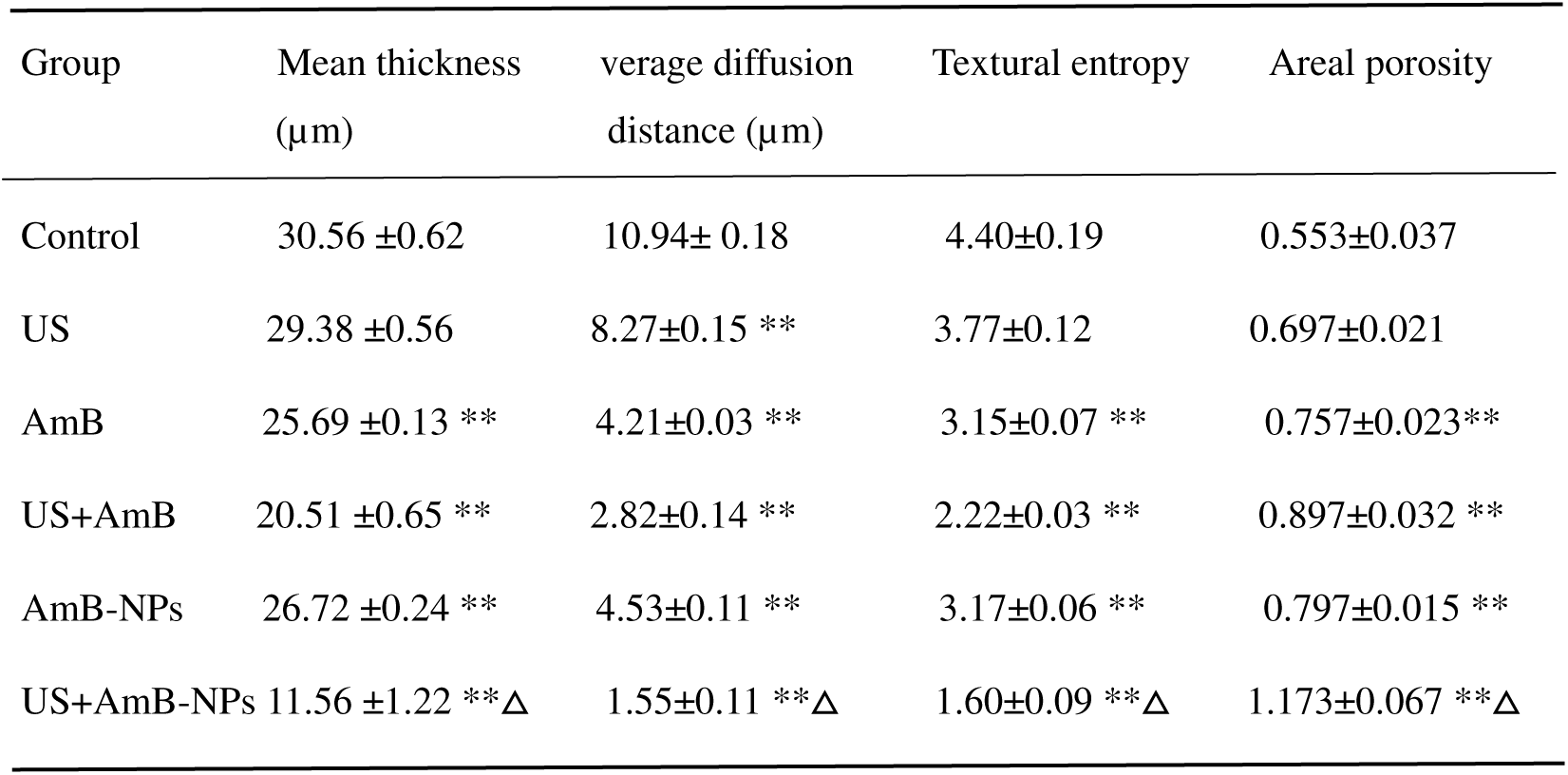
COMSTAT analysis of structural parameters of *C. albicans* biofilm in different treatments.

### Proteolytic and phospholipase enzymatic activities

The proteolytic and phospholipase enzymatic activities of *C. albicans* biofilm, which are both important virulence factors contributing to host tissue damage, were analyzed as shown in Fig. 6. When compared with the control group, the activity of protease and phospholipase in the combined ultrasound and AmB-NPs group was reduced by 68.41% and 68.57%, respectively, (P<0.01), while the enzyme activity was lower than that of the AmB group (P<0.05). These findings indicated that the combination of ultrasound and AmB-NPs could increase the ability to inhibit protease phospholipase activity, while weakening the expression of virulence factors, thereby reducing the pathogenicity of biofilms.

**Fig. 6.**
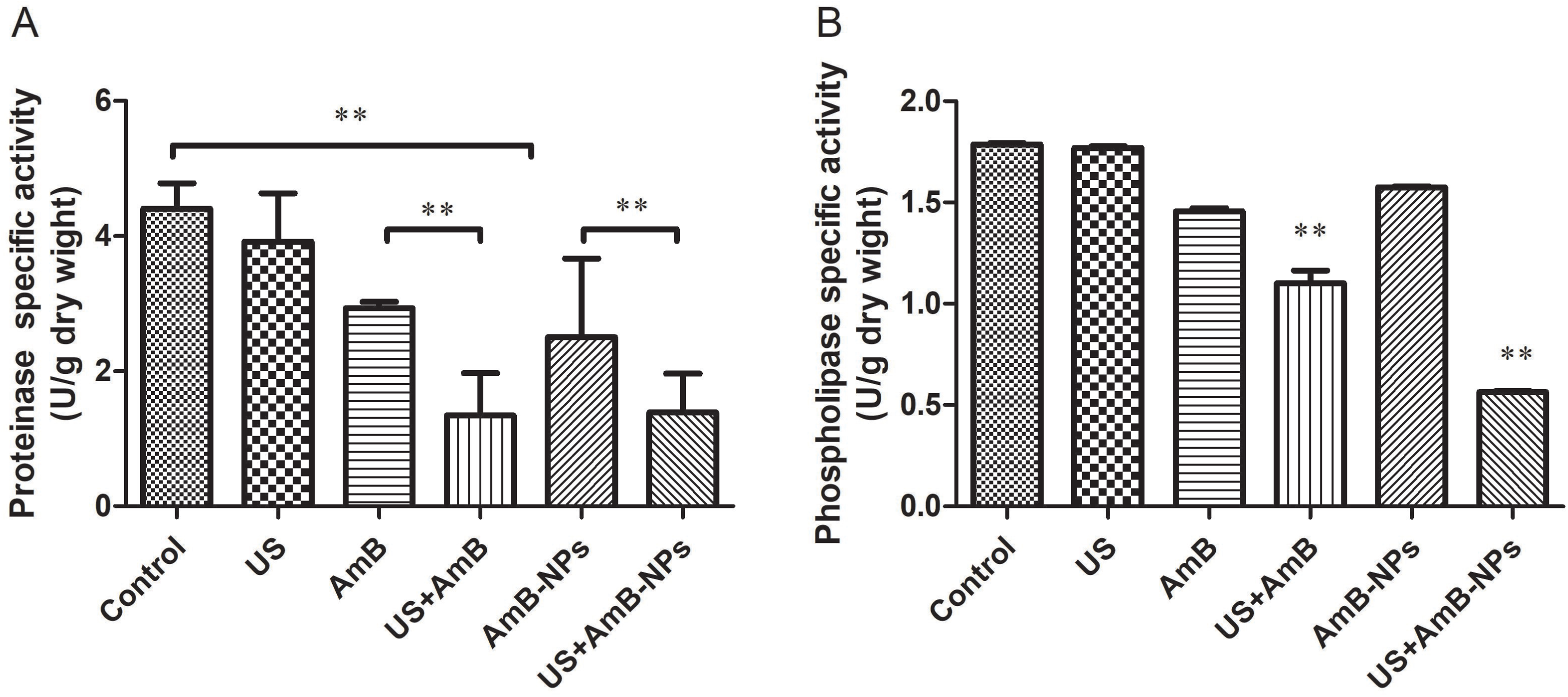
The proteolytic (A) and phospholipase (B) enzymatic activities of *C. albicans* biofilm of different treatment groups. Significant reductions in proteinase and phospholipase enzyme activities were observed in the combination treatment at 4.0 μg/mL AmB. The double asterisk (^∗∗^) denotes a statistically significant difference (P<0.01). US: ultrasound; AmB: amphotericin B; AmB-NPs: amphotericin B-loaded PLGA nanoparticles.

### Synergistic antifungal activity in vivo

The fungus loading onto the implantation catheter was quantified by CFUs after the corresponding treatment for 3 or 7 days and the colonies incubated on the plate are shown in Fig. 7. The colony counts were obviously reduced on the plates following treatment with ultrasound combined with AmB-NPs (Fig. 7A). Moreover, both free AmB and AmB-NPs could reduce the amount of fungus on the catheter. However, the fungal burden of rats treated with ultrasound combined with AmB-NPs was significantly lower than that of free AmB-treated rats or those that received other treatments (P<0.01), while ultrasound treatment alone did not significantly decrease the fungal counts compared with the control group (P > 0.05). The catheter fungal load showed a similar decrease trend after the 7-day-treatment experiment, with a significantly decline being observed between the 3 and 7-day-treatments following combined treatment with ultrasound and AmB-NPs (P<0.01) (Fig.7B). These results suggest a remarkably synergistically antifungal effect of ultrasound combined with AmB-NPs on C. albicans biofilm relative to AmB alone in vivo, which was in accordance with the results observed in vitro.

**Fig. 7.**
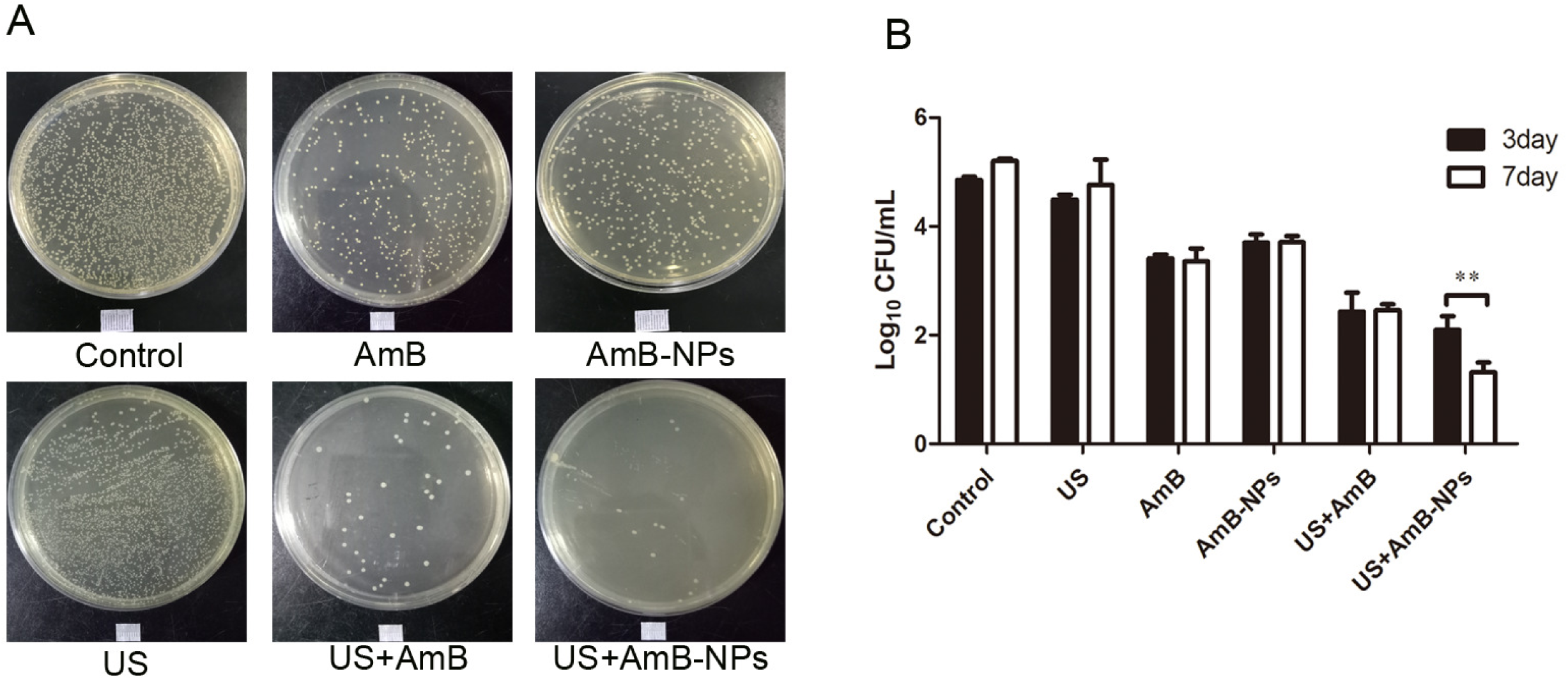
Catheter fungus loading of *C. albicans* with different treatments after 24 h of incubation on plate (A) and fungal colony counts (Log_10_ CFU) after 3 or 7 days of treatment (B). Double asterisk (^∗∗^) denotes a significant difference (P < 0.01).The bar indicates 50 μm. US: ultrasound; AmB: amphotericin B; AmB-NPs: amphotericin B-loaded PLGA nanoparticles.

To further observe the morphological changes in catheter biofilms after different treatments, the explanted catheters were stained with ConA and observed under CLSM. Figure 8 shows a mature biofilm architecture with abundant fungal cells embedded within the extracellular matrix on the catheter surface in the controls. Following treatment with AmB with or without ultrasound treatment, the biofilm structure was destroyed, the extracellular matrix was reduced, mycelia were shortened, and only a small amount of biofilm remained on the surface of the catheter. However, after the joint effects of ultrasound with AmB-NPs, only a single fungal colony remained on the catheter surface and the biofilm was substantially eliminated.

**Fig. 8.**
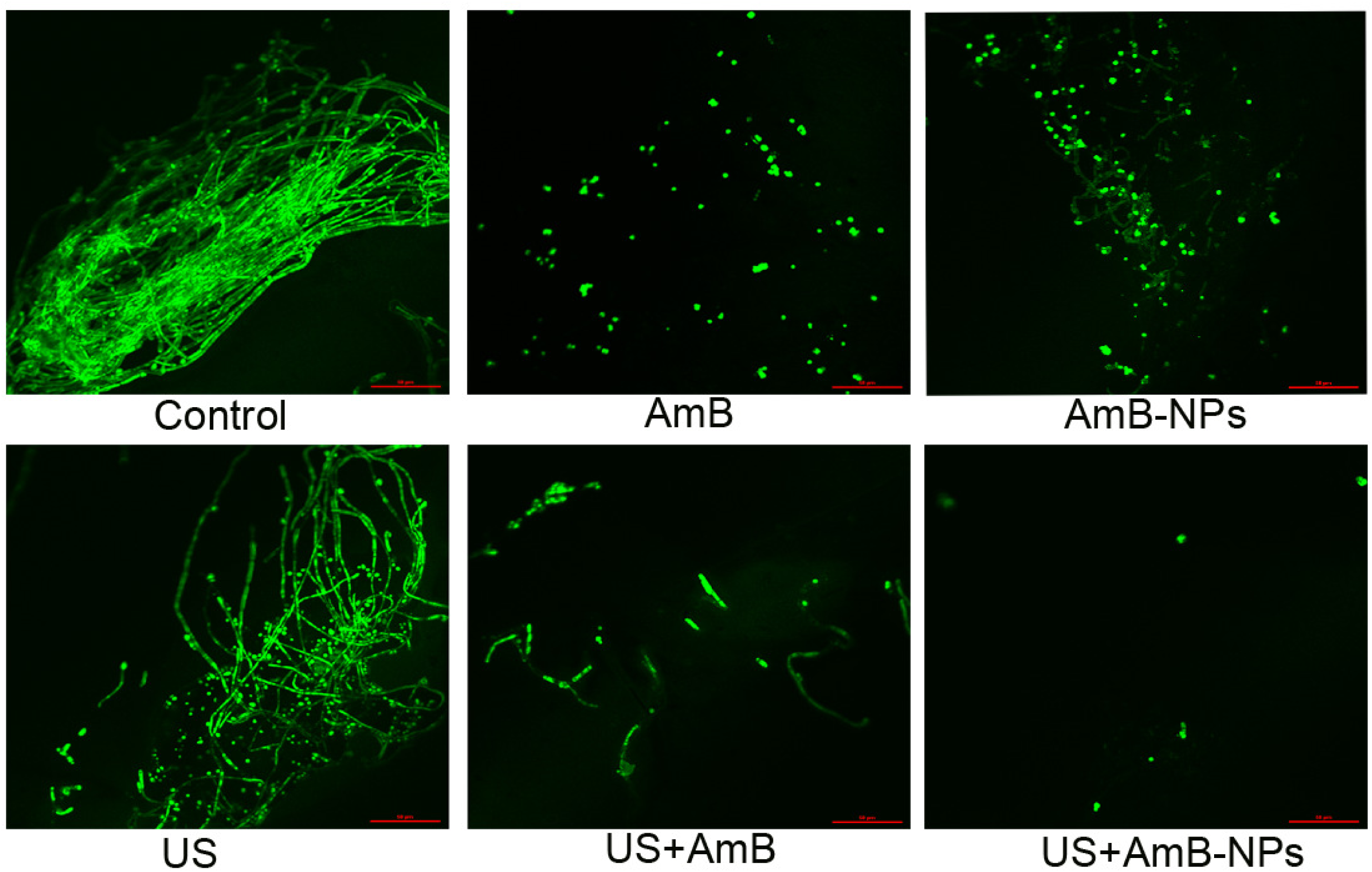
Morphological changes in catheter biofilm after seven-consecutive days of treatment. *C. albicans* cells were stained with ConA and visualized by CLSM at 400× magnification. The control group of mature biofilms with a dense structure on the catheter surface. In the group treated with ultrasound and AmB-NPs, only a single fungal colony remained on the catheter surface and the biofilm was substantially eliminated. The bar indicates 50 μm. US: ultrasound; AmB: amphotericin B; AmB-NPs: amphotericin B-loaded PLGA nanoparticles.

## Discussion

Fungal colonization of medical devices is a widespread problem and is responsible for nosocomial infections. *C. albicans* biofilm formation, which is commonly associated with implanted medical devices, is less susceptible to antifungal agents than other fungi and is therefore difficult to cure^[20]^. AmB has been used for decades to treat common invasive fungal infections. Although AmB has high nephrotoxicity, nanoparticle formulations of AmB have been shown to have reduced toxicity compared to its monomer state ^[21-22]^. Similarly, the results of the present study revealed lower toxicity toward red blood cells’ hemolytic activity and macrophage viability of AmB-loaded PLGA nanoparticles compared to free AmB (Fig. 3), which was associated with drug encapsulation and its subsequent slow release from nanoparticles. In vitro US-triggered drug release experiments showed that the release of AmB-NPs was delayed when allowed to proceed naturally but was significantly increased after sonication. These findings indicate that ultrasound promotes the diffusion of nanoparticles within the shells, resulting in increased drug release rates, which may realize the targeted release of drug-loaded nanoparticles at specific positions in which the drug vehicle receives ultrasound energy^[23-24]^.

Use of low-frequency ultrasound (20-100 KHz) as a non-invasive treatment has been shown to improve cell membrane permeability, increase drug penetration and achieve synergistic antibacterial effects by previous researchers^[25-26]^. The non-injury threshold of percutaneous irradiation has been verified to be under 0.30 W/cm2 for 15 min in our previous experiments^[19]^, and this ultrasound exposure dose is below the damage threshold reported by Schneider based on neuron loss and death of brain cells of continuous exposure at 2.6 W/cm2 for 20 min at 20 kHz^[27]^.

In the in vitro synergistic antifungal efficacy experiments conducted in this study, we quantitatively and qualitatively analyzed the metabolic activity and structural changes in biofilms by the XTT reduction assay and CLSM images taken after the application of different treatments. The results showed that joint treatment with ultrasound irradiation and AmB-NPs not only significantly reduced the activity of biofilm fungal cells (Fig. 4A) but also had a destructive effect on the structure of the biofilm itself, including reducing the thickness, blocking the supply of nutrients to the biofilms and rarefying the structure (Table 2). The highly effective synergistic antibacterial effect of low-frequency ultrasound combined with drug-loaded nanoparticles may be due to acoustic cavitation^[28]^. Specifically, the cavitation effect with the accompanying extreme physical phenomena, such as high pressure and high shear force, can loosen the densified structure of the biofilm extracellular matrix, thereby enhancing the drug permeability of the biofilm. Moreover, the shock waves and micro-jets generated by the ultrasonic cavitation effect produce sonoporation on the bacterial cell membrane interface. This temporary channel likely increases the amount of antibacterial drugs entering the cells. Moreover, drug-loaded nanoparticles as exogenous systemic administration of microbubbles greatly increased the number of cavitation nuclei, reduced the threshold of localized cavitation and enhanced the cavitation effect; therefore, ultrasound-irradiated nanoparticles break and release drugs to exert a highly effective antibacterial effect^[29-31]^.

In the in vivo synergistic antifungal efficacy experiments, we established an animal model of a *C. albicans* biofilm using the subcutaneous implantation of small pieces of biomaterials infected with *C. albicans* cells, and the mature biofilms were recovered two days after implantation. The infected rats were then treated with free AmB or AmB-NPs and ultrasound separately or together for 7 days. There was a remarkable reduction in the number of CFUs of the catheter fungus observed in the ultrasound combined with AmB-NPs treatment relative to AmB alone or ultrasound combined with AmB after 7 days of treatment (Fig. 7). Furthermore, CLSM images of the catheter biofilm after the different treatments were consistent with the antifungal effects of catheter fungus-loading. Specifically, the images revealed that the attached biofilm structure on the catheter was destroyed, the mycelium had disappeared, and only a few colonies remained (Fig. 8). These findings indicated that combined treatment with ultrasound and drug-loaded nanoparticles exert synergistic antifungal activity and achieve comparable or even better therapeutic effects at lower doses, which reduces the incidence of adverse effects. Although treatment with a combination of AmB and echinocandin or azoles produced good results in vitro and in animal models of invasive fungal infection, the combination of drugs will often increase the side effects of drugs to varying degrees and the clinical treatment effect is not satisfactory^[32-33]^. Thus, ultrasound may be a potential way for inducing synergistic antibacterial effects when compared to traditional combination therapy.

In summary, we prepared AmB-NPs that slowly released the drug, resulting in lower toxicity compared to free AmB. And more importantly, the synergistic antifungal efficacy of low-frequency and low-intensity ultrasound combined with AmB-loaded PLGA nanoparticles on *C. albicans* biofilms was successfully supported in vitro and in vivo assays. The combination of ultrasound with drug-loaded nanoparticles can be considered one promising strategy to improve efficacy, reduce dose of west medicine and shortening the therapeutic period for antifungal therapy. Future perspectives include the potential to use ultrasound with drug-loaded nanoparticles to treat common clinical *C. albicans* biofilms-related infections, such as those associated with central venous catheter insertion, catheters and artificial joint replacement.

## Materials and methods

### Materials

AmB (99.8%purity), polyvinyl alcohol (PVA), 2,3-bis(2-methoxy-4-nitro-5-sulfo-phenyl)-2H-tetrazolium-5 -carboxanilide(XTT), menaquinone, 3-(4-5-dimethylthiazol-2-yl)-2,5-diphenyl tetrazolium bromide (MTT), crystal violet, concanavalin A (Con A), azocasein and phosphatidylcholine substrate and dialysis bags (molecular weight cut-off of 12 kDa) were purchased form Sigma-Aldrich (St. Louis, MO, USA). PLGA polymer material with a molecular weight of 21kDa (ratio of lactide to glycolic acid molar ratio of 50:50) was purchased from RuiJia Biological (XiAn, Chain). RPMI 1640 medium, fetal bovine serum (FBS) and phosphate-buffered saline (PBS) was purchased from Gibco BRL (Carlsbad, CA, USA). Sabouraud Dextrose Agar (SDA) and Sabouraud Dextrose broth (SD) were obtained from Huankai Microbial Co., Guangdong, China. A LIVE/DEAD BacLightTM Bacterial Viability Kit was acquired from Invitrogen (L7012, Invitrogen, CA, USA).

### *C. albicans* strains and biofilm formation

*C. albicans* ATCC10231 was obtained from the China General Microbiological Culture Collection Center. Individual colonies were selected from the SDA plate and inoculated into 100 mL of SD broth at 37°C for 24 h with agitation (150 rpm) to prepare *C. albicans* suspensions, which were preserved at 4°C for future experiments.

For biofilm formation, we developed a model of fungal biofilm growth of *C. albicans* by continuous incubation for 48 h on a 35 mm diameter plastic bottom petri dish in vitro (supplementary material 1). Biofilm formation was quantified by crystal violet staining and XTT reduction assay^[34]^.

### Animal species

Sixty healthy female Sprague-Dawley (SD) rats weighing about 200 g were obtained from the Laboratory Animal Administration Center of Chongqing Medicine University. All animals were housed in monomer independent air cages (IVC) and given standard diet ad libitum. The animal experiments conducted in this study were performed with the approval of the Chongqing Medical University Institutional Animal Care and Use Committee.

### Preparation and characterization of AmB-loaded nanoparticles

AmB-loaded PLGA nanoparticles (AmB-NPs) and plain PLGA nanoparticles (NPs) were formulated by a double emulsification method as previously described^[19]^. Briefly, a preweighed amount of AmB powder was dissolved in DMSO that were miscible with water (20:80%, v/v). PLGA polymer material was dissolved completely in dichloromethane and mixed with the drug or water for the first ultrasonic oscillation (XL2020, USA) at 100W for 2 min. Next, 1% PVA aqueous solution was added to the polymeric mixture for the second ultrasonic oscillation at 100W for 5 min. NPs were prepared following a similar method, except that the drug was exchanged for an equal amount of deionized water. The nanoparticles were resuspended, washed and collected after the organic solvent was evaporated at room temperature for 6 hours. The physical characteristics of nanoparticles, including the average diameter (D), zeta potential (ZP) and polydispersity index (PDI), were measured using a Malvern laser particle size analyzer (Zeta SIZER 3000HS, USA) and morphological characterization of nanoparticles was conducted by transmission electron microscopy ((TEM, Hitachi High-Technologies, Tokyo, Japan). The drug loading content (LC%) and the encapsulation efficiency (EE%) of AmB-NPs were then calculated using the following equations:

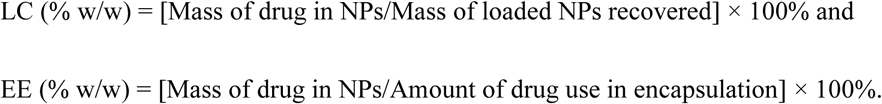

### In vitro investigations of ultrasound-triggered drug release

The kinetic release of AmB from nanoparticles in vitro with ultrasonication was assessed. A sample of AmB-NPs lyophilized powder was diluted in PBS with or without sonication (fixed frequency of 42 kHz) at an intensity of 0.30 W/cm^2^ for 15 min. After sonication, the samples were then individually transferred into dialysis bags, which were subsequently immersed in a container filled with 50 mL PBS and shaken at 100 rpm. Dialysate (1 mL) samples were collected and the percentage of AmB released from nanoparticles was evaluated by UV-vis spectrophotometry (UV-2600 SHIMADZU, Japan) at each predetermined time point. Samples of AmB-NPs that underwent the same procedure, but without sonication, were used as controls.

### Hemolytic activity of AmB-NPs in vitro

AmB sometimes causes significant adverse effects such as hematological toxic reactions, which induce hemolytic anemia by allowing normal red blood cells to undergo hemolysis^[7]^. The hemolytic activity of AmB-NPs was assessed using a previously reported method^[35]^. Briefly, 1.0 mL of 1% healthy rabbit red cell suspension was mixed with 1.0 mL of free AmB, AmB-NPs or NPs solution, and final AmB concentrations of 0–32 μg/mL were applied in the free AmB and AmB-NPs group, respectively. The corresponding concentrations of NPs were 0–800 μg/mL (based on drug loading content). After 24 h of incubation, the supernatants were collected and the absorbance at 540 nm was measured using a spectrophotometer. Additionally, red cells incubated with PBS alone served as a negative control group (estimate the natural hemolysis) and those incubated with distilled water as a positive control (serve as 100% hemolysis). The percentage of hemolysis induced by free AmB or AmB-NPs was then calculated using the following equation:

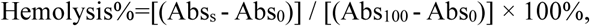

where AbsS is the average absorbance of the sample, Abso is the average absorbance of the negative control, and Abs100 is the average absorbance of the positive control.

### Effects of AmB-NPs on macrophage activity

Macrophages are major anti-inflammatory cells that play an important role in eliminating extraneous pathogens and participate in immune responses^[36]^; therefore, it is necessary to investigate the effects of AmB-NPs on macrophage activity. The free AmB, AmB-NPs, and NPs were co-cultured with RAW264.7 macrophages in the logarithmic phase for 24 h, and the final concentration of the drug used in this experiment was 4.0 μg/mL (based on a test of the susceptibility of *C. albicans* biofilms to AmB). NPs at corresponding concentrations (100 μg/mL calculated by drug loading content) were used in this experiment. Finally, the activity of macrophages in each group was detected by MTT assay based on the absorbance at 490 nm ^[37]^.

### Ultrasonic irradiation method in vitro and in vivo

The ultrasound device provided by Haina Science and Technology Co., Suzhou, China, was used in this study for in vitro and vivo experiments. The instrument parameters were a fixed frequency of 42 kHz, a transducer diameter of 5.0 cm and an adjustable sound intensity output of 0–0.6 W/cm^2^, which was monitored using an acoustic power detector UPM-DT-1AV (OHMIC Instruments, USA). The formula used to calculate the spatially and temporally averaged intensity (I_SATA_) (W/cm^2^) was equal to the ultrasonic power (W) divided by the transducer surface area (cm^2^)^[38]^.

For sonication in vitro, a biofilm growing plastic bottom petri dish was fixed in a degassed distilled water bath using an aluminum wire. The ultrasonic transducer was submersed directly on the bottom of a biofilm plate and degassed distilled water was used as the ultrasonic medium. The irradiation setup was submersed completely in a degassed distilled water bath and maintained at room temperature (22°C). For sonication in vivo, the medical ultrasonic coupling agent was uniformly coated on top of the transducer, after which the transducer was affixed tightly to the skin of the rat for irradiation after injection of the drug into the local infection site. A schematic diagram of the ultrasonic irradiation method is shown in Fig. 1.

### Synergistic effects of ultrasound combined with AmB-NPs on *C. albicans* biofilms in vitro

*C. albicans* biofilm in the mature period after 48 h incubation was observed microscopically and evaluated using a growth dynamics curve (Fig. S1). To further investigate whether there is a synergistic effect of ultrasound combined with AmB-NPs on *C. albicans* biofilm in vitro, the formation biofilm were performed by ultrasound and AmB-NPs separately or jointly in the following treatments: untreated (Control), ultrasound alone (US), free AmB alone (AmB), ultrasound combined with free AmB (US+AmB), AmB-loaded nanoparticles (AmB-NPs), and ultrasound combined with AmB-loaded nanoparticles (US+AmB-NPs). The susceptibility of AmB and AmB-NPs to *C. albicans* biofilms was determined and the concentration of sessile minimum inhibitory concentration (SMIC)_50_ of AmB was 4.0 μg/mL (Fig. S2). Therefore, the final concentration of AmB employed in the AmB group, AmB-NPs group, US+AmB group or US+AmB-NPs group was 4.0 μg/mL. Furthermore, the effects of ultrasound on *C. albicans* biofilm following different acoustic parameters showed that ultrasonic irradiation at 0.30 W/cm^2^ for 15 min may provide favorable conditions for synergistic effects of ultrasound combined with AmB-NPs (Fig.S3); therefore, the ultrasound irradiation in this experiment was performed at an intensity of 0.30 W/cm^2^ for 15 min. The biofilm activity after 24 h of incubation following treatment was measured by a XTT reduction assay. The absorbance reflects the activity of cells within the biofilm as measured in a microplate plate reader (ELX800, BIO-TEK Instruments, Inc.) at 490 nm. The total biomass of biofilm treatment was quantified according to the reported method using 0.1% crystal violet staining at 570 nm absorbance in a spectrophotometer^[39-40]^. All experiments were conducted independently and in triplicate.

For analysis of the biofilm structure changes after treatment, a LIVE/DEAD BacLightTM Bacterial Viability Kit was used to stain *C. albicans* biofilm, which is a dual dye specifically designed for rapid and accurate detection of fungal cell activity, including SYTO9 and PI dye. Following treatment of *C. albicans* biofilm by ultrasound and AmB-NPs separately or jointly, 50 μL SYTO9/PI fluorescent dye was added to the biofilm petri dish and incubated at 37°C for 30 min in the dark. The morphology and distribution of live and dead fungal cells of biofilm were then visualized by confocal laser scanning microscopy (CLSM, A1+R, Nikon, Tokyo, Japan). Next, biofilm in the field from three randomly selected positions was scanned from the bottom layer to the top layer at a layer spacing of 0.5 μm and the three-dimensional (3-D) structural image of the biofilm was reconstructed. The structure of *C. albicans* biofilms was then assessed using the COMSTAT software (presented by Professor Haluk Beyenal of Montana State University, USA) to determine the mean thickness (μm), textural entropy (TE, reflects biofilm heterogeneity), areal porosity (AP, reflects biofilm gap channels) and average diffusion distance (ADD, reflects biofilm nutrient supply distance)^[41]^.

### Proteinase and phospholipase enzyme secretion assay

Secretory acidic proteases and extracellular phospholipase are two important virulence factors of *C. albicans* that are closely related to their invasiveness and pathogenicity. To investigate whether ultrasound affected the enzyme secretion activity, proteinase and phospholipase enzyme secretion assays were conducted as previously described^[42-43]^. Briefly, after the biofilm was treated by ultrasound combined with AmB-NPs, it was rinsed repeatedly in a Petri dish, then suspended in PBS. The proteinase enzyme activity was subsequently determined by the supernatant of the biofilm solution mixed with 1% azocasein at 1:9 (v/v) and incubated for 1 hour at 37°C in 5% CO_2_. The reaction was stopped by adding 500 μL of 10% trichloroacetic acid, then incubated for another 10 min at room temperature after which it was centrifuged at 10000 rpm for 5 min. Next, 500 μL of supernatant was mixed with 500 μL of 0.5 M sodium hydroxide, which was incubated for 15 min at 37°C in 5% CO_2_. The absorbance of the supernatant was measured in a spectrophotometer at 440 nm. The phospholipase enzyme activity was determined by mixing the supernatant of the biofilm solution with an equal volume of phosphatidylcholine substrate for 1 hour at 37°C in 5% CO_2_, after which the absorbance at 630 nm was read in a spectrophotometer. Finally, the enzyme activity was reported as a specific activity unit (U) that increases the absorbance by 0.001 per minute and normalized by the dry weight of biofilms (U/g dry weight).

### Subcutaneous catheter of biofilm formation in a rat model and antifungal therapy

The method of establishing an subcutaneous catheter of biofilm formation in an in vivo rat model was conducted as described by Ricicová et al.^[44]^ as described in detail in the supplementary material. The intravenous catheters (diameter of 2.4 mm) were cut into segments about 1.0 cm in length, and then were adhered fungus colonies via co-incubation with *C. albicans* suspension (10^7^ CFU/mL) for 3 h at 37°C and 120 rpm. The catheters were subsequently implanted under the skin of SD rats. At 48 hours after implantation, adhesion of fungus colonies onto the catheter reached a stable period and biofilm was formed on the surface of the subcutaneous catheter, which was confirmed by calculation of catheter fungus loading and microscopic observation (Fig. S4).

To further investigate the synergistic antifungal efficacy of ultrasound mediated AmB-NPs on *C. albicans* biofilms in vivo, the infected rats were randomly divided into the following six groups: untreated (Control), ultrasound alone (US), free AmB alone (AmB), ultrasound combined with free AmB (US+AmB), AmB-loaded nanoparticles (AmB-NPs), and ultrasound combined with AmB-loaded nanoparticles (US+AmB-NPs). The dose of free AmB was selected based on recommendations for the clinical treatment of fungal infection and converted to the body weight of rats, which was 1 mg/kg/day for 7 days (total of 7.0 mg/kg/animal). The dose of AmB-NPs at corresponding concentrations was 10 mg/kg/day for 7 days, totaling 70 mg/kg/animal. The animals in some groups were subjected to once-daily ultrasonic irradiation for 15 min at 0.30 W/cm^2^ with a continuous wave. This level of treatment was previous shown not to cause skin lesions during bio-safety assessment of percutaneous irradiation in our previous experiments^[19]^. The infected rats were divided into two groups for 3 day and 7 day treatments, respectively. After the third and seventh days of continuous treatment, the explanted catheters were removed (at 24 h after last treatment) and biofilms were subsequently stripped from the catheter surface and dispersed in PBS by sonication for 5 min at 40 kHz in a water bath sonicator (KQ5200DE, SuZhou, China). Our preliminary results indicated that this process did not affect cell culturability. Samples were then seeded on SDA plates by 1:10 dilution for 24 h at 37°C, after which the number of CFUs on the catheters were counted. For observation of morphological changes in the catheter biofilm, explanted catheters were stained with 50 μg/mL ConA (which binds to polysaccharides to outline the cell walls of the yeast and produce intense green fluorescence) for 1 hour in the dark. Fluorescence images were observed under CLSM at excitation/emission wavelengths of 488/530 nm.

### Statistical analysis

Qualitative data in this study were described as the means ± standard deviation (X ± SD) of three experiments and analyzed using GraphPad Prism version 6.00 for Windows (GraphPad Software; La Jolla, CA, USA). Results were considered to be significant when p-values were <0.05.

## Acknowledgments

This research was funded by the Chongqing Research Program of Basic Research and Frontier Technology (No. csct2016jcyjA0098), Program of Chongqing Special Social Livelihood of the People of Science and Technology Innovation (Nocstc2016shmszx130029). In addition, we would like to thank LetPub (www.letpub.com) for providing linguistic assistance during the preparation of this manuscript.

## Conflict of Interests

The authors declare, to the best of their knowledge, that there is no conflict of interest regarding the publication of this paper.

---

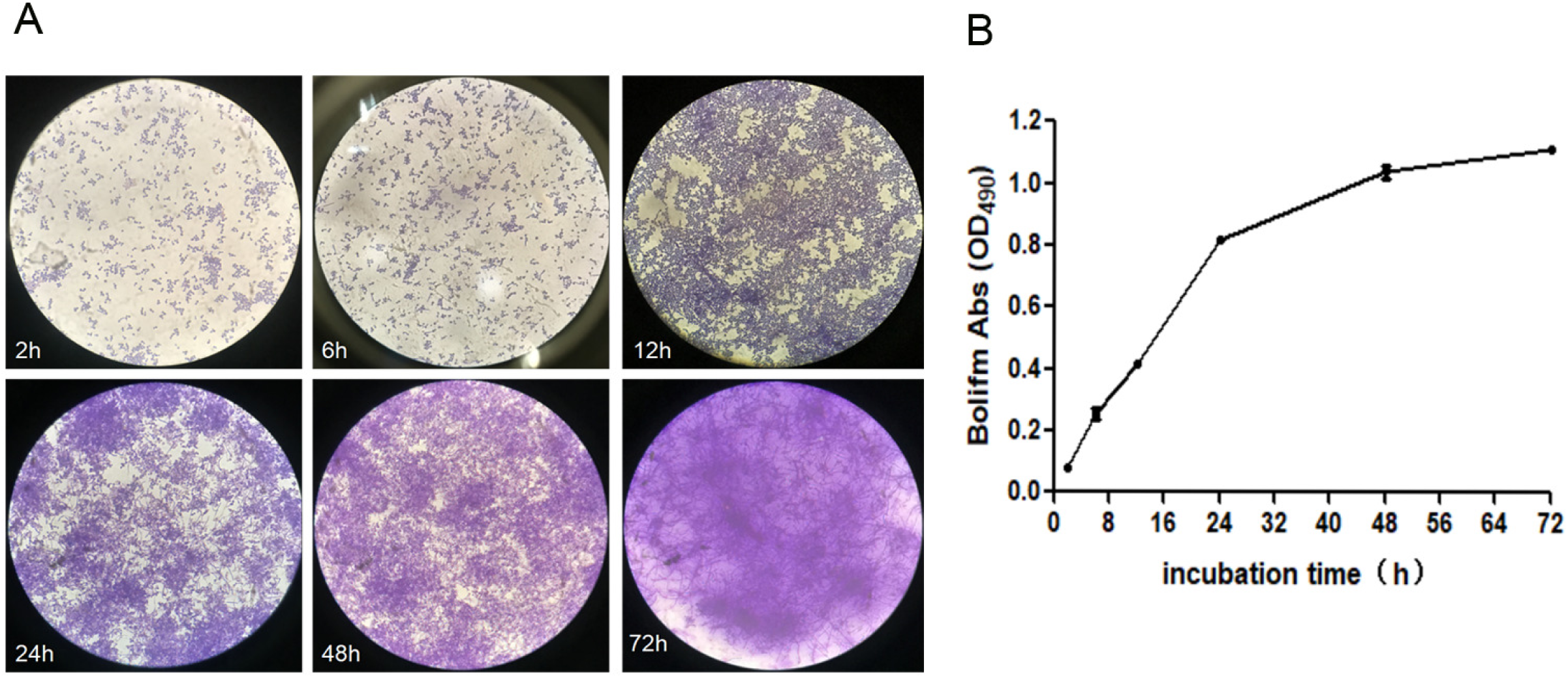

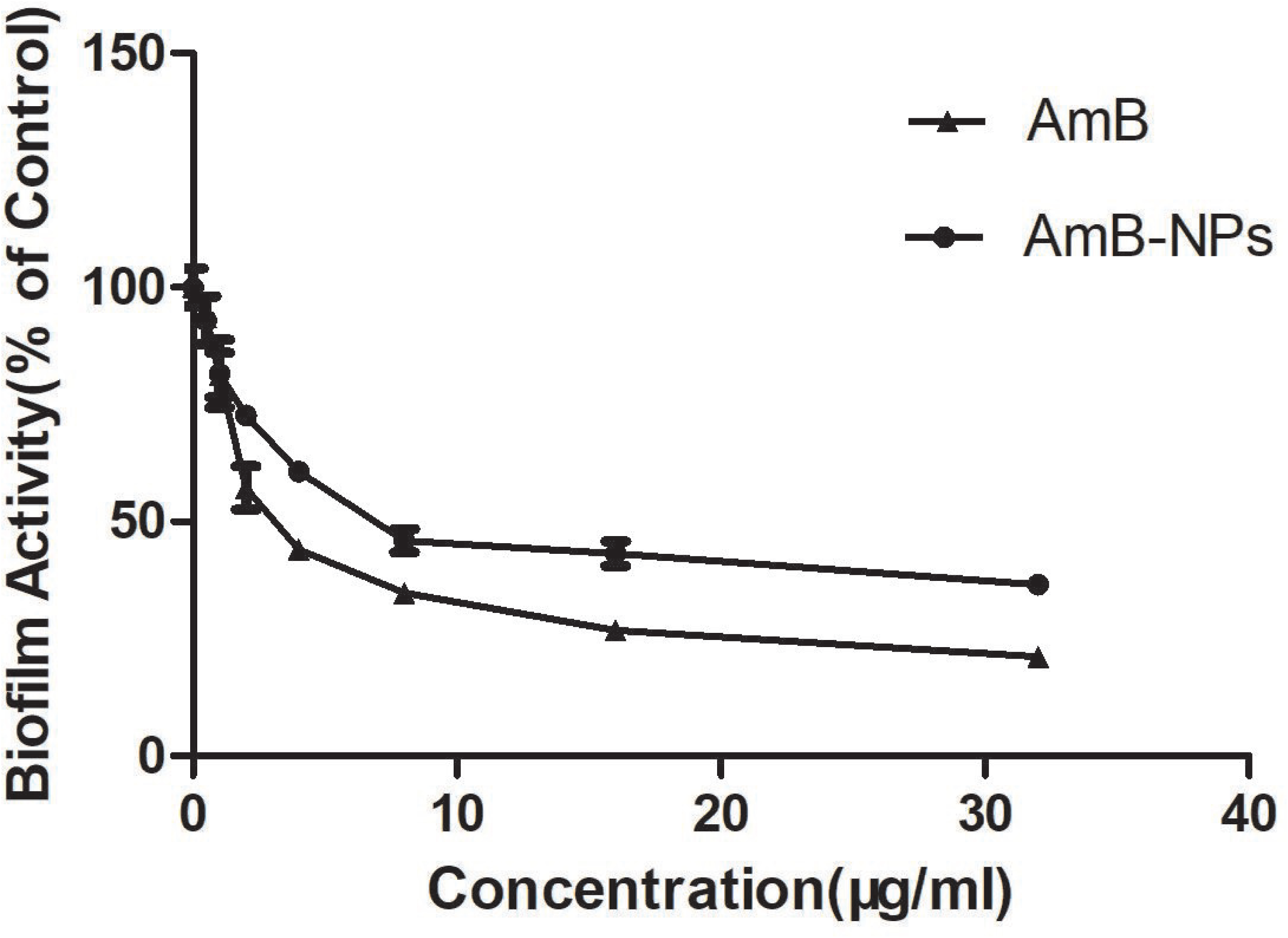

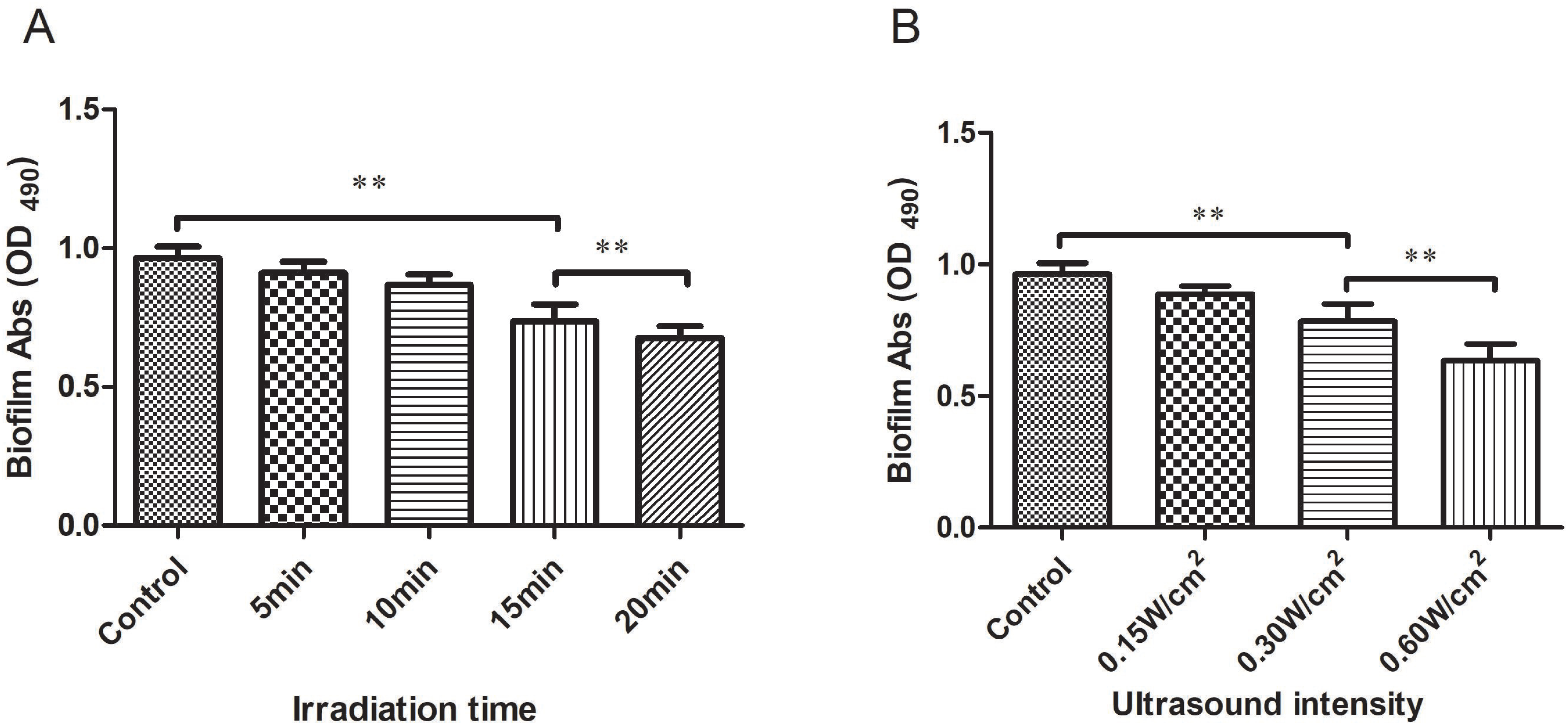

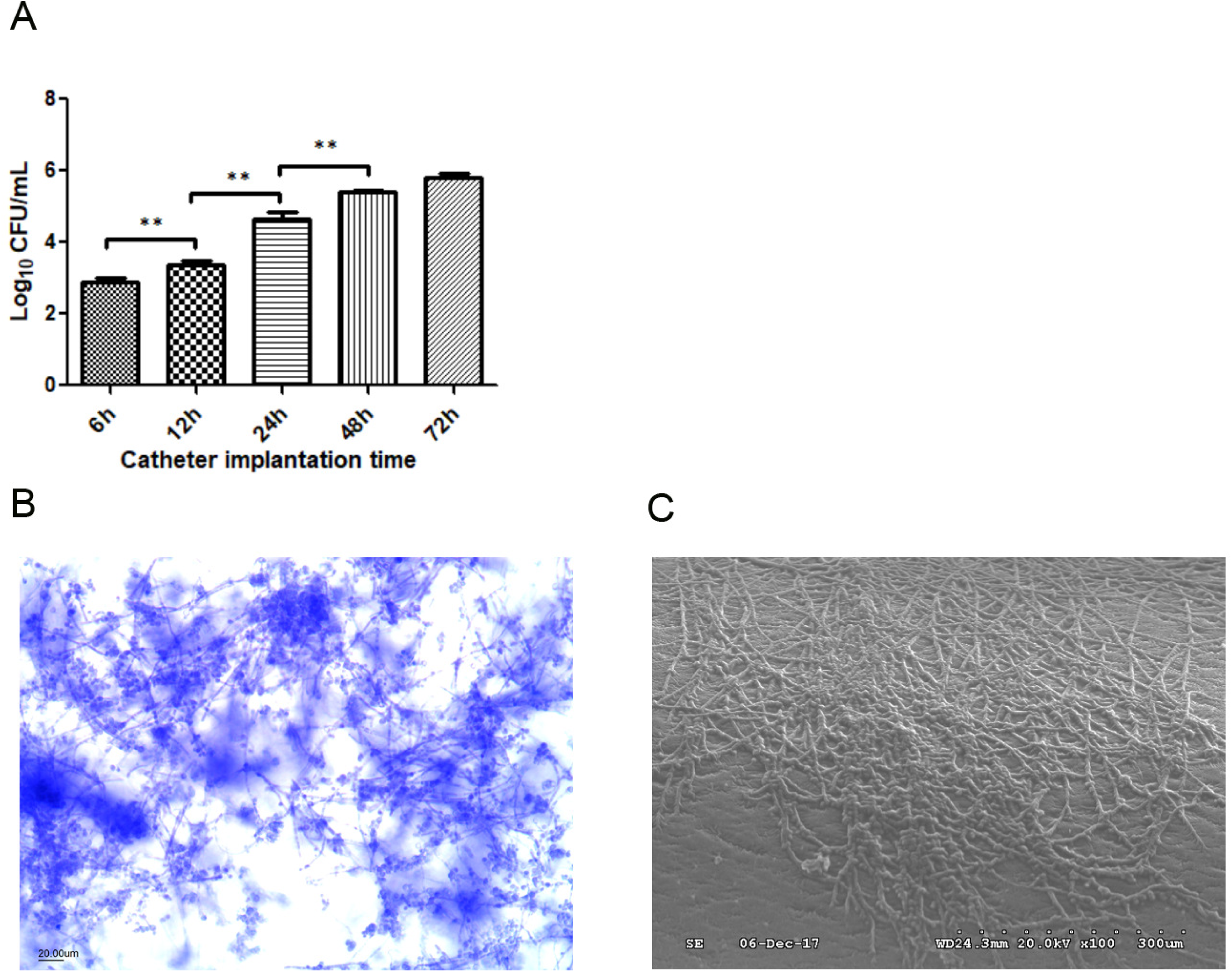

